# Prelamin A Does Not Promote Atherosclerosis or Vascular Smooth Muscle Loss

**DOI:** 10.1101/2025.09.15.676432

**Authors:** Yuexia Wang, Leroy C. Joseph, Cecilia Östlund, George Kuriakose, Wei Hsu, Susan Michaelis, Howard J. Worman

## Abstract

**BACKGROUND:** Hutchinson-Gilford progeria syndrome (HGPS) is an accelerated aging disorder characterized by numerous symptoms, including early-onset atherosclerosis, with most patients suffering fatal myocardial infarctions or strokes by the second decade of life. HGPS is caused by mutations in *LMNA* that lead to expression of an internally truncated, farnesylated prelamin A variant called progerin, which induces loss of vascular smooth muscle cells (VSMCs). Some studies have also reported that accumulation of full-length farnesylated prelamin A, which is normally completely processed to mature non-farnesylated lamin A, can also drive vascular pathology during physiological aging.

**METHODS:** To assess the effects of prelamin A expression on atherosclerosis and aortic VSMCs, we used *Lmna*^L648R/L648R^ mice that express a prelamin A variant with a lysine to arginine point mutation that prevents its processing to mature lamin A. To determine if prelamin A expression has an impact on atherosclerotic plaques, we crossed *Lmna*^L648R/L648R^ mice to LDL receptor-deficient *Ldlr*^−/−^ mice that develop hyperlipidemia on a high-fat diet.

**RESULTS:** Atherosclerotic plaque lesion area and necrotic core area were not different in hyperlipidemic *Lmna*^L648R/L648R^ mice that expressed only prelamin A, and no mature lamin A, compared to hyperlipidemic *Lmna^+/+^* mice that expressed only fully-processed mature lamin A and no prelamin A. Additionally, exclusive prelamin A expression did not result in loss of aortic VSMCs or adventitial thickening in hyperlipidemic *Lmna*^L648R/L648R^ mice with atherosclerosis at 28 weeks of age. Indeed, aortic vascular smooth muscle remained normal in older *Lmna*^L648R/L648R^ mice at 52 weeks of age.

**CONCLUSIONS:** In contrast to the prelamin A variant progerin expressed in HGPS, prelamin A does not appear to cause vascular smooth muscle loss, promote atherosclerosis or drive vascular aging.

## INTRODUCTION

The nuclear lamina is a meshwork of intermediate filaments on the inner aspect of the inner nuclear membrane.^1^ The lamin A/C gene (*LMNA*) encodes prelamin A and lamin C, two of the intermediate filament protein building blocks of the lamina.^2–4^ Prelamin A, but not lamin C, has a carboxyl-terminal cysteine-aliphatic-aliphatic-any amino acid (CAAX) motif that initiates a series of posttranslational processing reactions to generate mature lamin A.^5,6^ After the CAAX processing reactions of farnesylation of the cysteine, cleavage of -AAX, and carboxymethylation of the cysteine, prelamin A undergoes a final cleavage reaction catalyzed by the zinc metalloprotease ZMPSTE24. This removes the last 15 amino acids of prelamin A, including its farnesylated cysteine, resulting in the production of mature, unfarnesylated lamin A. Genetic mutations leading to defects in the ZMPSTE24-catalyzed cleavage of prelamin A cause disorders with features of accelerated aging.

Hutchinson–Gilford progeria syndrome (HGPS) is an accelerated aging disorder caused by a splicing mutation in *LMNA* that generates a prelamin A variant with an internal in-frame deletion of 50 amino acids called progerin.^7,8^ Progerin retains its CAAX motif, but lacks the ZMPSTE24 cleavage site, and thus the carboxyl-terminal cysteine remains permanently farnesylated and methylated. Children with HGPS suffer from early-onset cardiovascular disease with myocardial infarction or stroke typically the cause of death in the mid-teens.^9^ Autopsy findings in HGPS have shown severe loss of vascular smooth muscle cells (VSMCs) in the atherosclerotic aortic media, along with adventitial fibrosis and thickening.^10,11^ Similarly, mouse models of HGPS that express progerin exhibit loss of media VSMCs and adventitial fibrosis and thickening.^12–16^ Crossing HGPS mice onto an *Apoe*^−/−^ or *Ldl*r^−/−^ background and feeding a high-fat diet also accelerates atherosclerosis accompanied by loss of VSMCs and adventitial thickening.^17,18^

Mandibuloacral dysplasia with type B lipodystrophy (MAD-B) and restrictive dermopathy are rare progeroid disorder caused by mutations in *ZMPSTE24*, leading to loss of enzyme activity and accumulation of full-length farnesylated prelamin A.^19–23^ MAD-B is characterized by growth retardation, partial lipodystrophy, mottled pigmentation of the skin and prominent bone defects including mandibular and clavicular hypoplasia, acroosteolysis, and delayed closure of cranial sutures.^19–23^ The greater the loss of ZMPSTE24 activity, the more severe the signs and symptoms, with total loss of function causing the neonatal lethal disorder restrictive dermopathy.^24,25^ While skeletal and cutaneous manifestations of MAD-B and HGPS overlap, there is no documentation of accelerated atherosclerosis in patients with MAD-B.^19–22,26^ Similarly, *Zmpste24*^−/−^ mice that accumulate prelamin A have overlapping phenotypes with HGPS model mice but do not have depletion of aortic VSMCs. However, *Zmpste24*^−/−^ mice have a median survival of only approximately 20 weeks and aortic vascular smooth muscle has only been examined at a maximum age of 21 weeks.^27–30^ It is therefore possible that prelamin A accumulation could lead to vascular smooth muscle defects in older mice.

We previously generated a *Lmna*^L648R/L648R^ mouse, which models a patient with the analogous human mutation, *LMNA* L647R, whose relatively mild disease called MAD-B-like, resembles MAD-B.^31,32^ The *Lmn*a L648R amino acid substitution abolishes the ZMPSTE24 cleavage site in prelamin A. *Lmn*a^L648R/L648R^ mice express only prelamin A (with a single amino acid substitution) and no mature lamin A. These mice have failure to thrive and bone defects similar to, albeit less severe and of later onset, than those in *Zmpste24*^−/−^ mice. In contrast to *Zmpste24*^−/−^ mice, which die by 4-7 months of age^27,28^, *Lmna*^L648R/L648R^ mice have near-normal longevity.^31^ Their less severe phenotype and longer lifespan allow for the analysis of how prelamin A impacts the vasculature and atherosclerosis at older ages.

A careful examination of the effects of prelamin A on vascular smooth muscle in older animals and in the setting of atherosclerosis is critical, considering previously reported findings suggesting that prelamin A may drive vascular disease in normal physiological aging. Ragnauth et al.^33^ reported that prelamin A accumulated in carotid and aortic media of aged but not young individuals and in atherosclerotic lesions, where it often colocalized with senescent and degenerating VSMCs. They further reported that ZMPSTE24 expression decreased in late-passage VSMCs, correlating with an increase in prelamin A, and that prelamin A induced senescence and disrupted mitosis in cultured VSMCs. In addition, Liu *et al*.^34^ presented findings that prelamin A accumulated in arteries from young children on dialysis who develop hallmarks of premature vascular aging, including arterial stiffening, loss of VSMCs, and medial vascular calcification.

To determine if prelamin A promotes vascular smooth muscle loss or has an impact on the severity of atherosclerotic plaques, we crossed *Lmna*^L648R/L648R^ mice to LDL receptor-deficient *Ldlr*^−/−^ mice that develop hyperlipidemia on a high-fat diet to determine whether hyperlipidemic stress might reveal a heretofore undetected vascular phenotype in *Lmna*^L648R/L648R^ mice. Our comparison of these hyperlipidemic *Lmna*^L648R/L648R^ and *Lmna*^+/+^ mice, however, suggests that prelamin A does not appear to cause vascular smooth muscle loss, nor promote atherosclerosis, nor drive vascular aging. Furthermore, non-hyperlipidemic *Lmna*^L648R/L648R^ mice demonstrate no vascular pathology at one year of age. These results are in contrast to the prelamin A variant progerin, expressed in HGPS, which drives vascular defects.^12–16^

## METHODS

### Data Availability

The data that support the findings of this study will be shared on reasonable request to the corresponding authors.

### Animals and Tissue Preparation

The Institutional Animal Care and Use Committee at Columbia University Irving Medical Center approved all protocols. We previously described the generation and characterization of *Lmna* L648R mice.^31^ Dr. Alan R. Tall (Columbia University) provided LDL receptor-deficient *Ldlr*^−/−^ mice, which have been descrbed previously.^35^ All mice were on the C57BL/6J genetic background. We performed PCR on genomic DNA isolated from tail clippings for genotyping. Mice were housed in a barrier facility with 12/12 h light/dark cycles. We fed mice a regular chow diet with 4.5% kcal from fat (5053-PicoLab® Rodent Diet 20, LabDiet), with some mice switching to a high-fat diet with 42% kcal from fat (TD.88137, Inotiv) for 12 weeks starting at 16 weeks of age. Mice were routinely weighed every four weeks, and once per week while receiving a high-fat diet. To estimate mean high-fat diet consumption, food was weighted before placing in cages and the left-over was measured when changing and adding new food weekly.

To obtain aortic roots for analysis of atherosclerotic lesions, we fed 16-week-old male mice a high-fat diet for 12 weeks. The 28-week-old mice were anesthetized with isoflurane and after collecting blood by left-ventricular puncture, the vasculature was perfused with cold phosphate-buffered saline. The heart with attached aortic root was fixed in 10% neutral buffered formaldehyde (Ricca Chemical), paraffin embedded and sectioned. The remainder of the aorta and liver were dissected and similarly fixed, embedded, sectioned, and stained with hematoxylin and eosin (H&E) and Masson’s trichrome. Aortas were also dissected from 52-week-old male mice and similarly prepared.

### Protein Extraction and Immunoblotting

Proteins were extracted from mouse liver and aorta tissue, separated by electrophoresis on SDS-polyacrylamide gels, transferred to nitrocellulose membranes and analyzed by immunoblotting using methods described previously.^36^ Primary antibodies used for immunoblotting were mouse anti-lamin A/C (Santa Cruz, #376248) at 1:1,000 dilution, rat monoclonal anti-prelamin A 3C8 (Sigma-Aldrich, MABT1458, clone 3C8) at 1:1,000 dilution, rabbit anti-phospho-AKT Ser473 (pAKT) (Cell Signaling, #4060) at 1:1,000 dilution, rabbit anti-AKT (Cell Signaling, #4691) at 1:1,000 dilution, and mouse anti-β-actin (Santa Cruz Biotechnology, sc-47778) at 1:4,000 dilution. Secondary antibodies were ECL-horseradish peroxidase-conjugated anti-rabbit, anti-rat, and anti-mouse antibodies (GE Healthcare) used at a dilution of 1:4,000. Signals were detected using SuperSignal West Pico PLUS Chemiluminescent Substrate (Thermo Fisher Scientific) and autoradiography film (LabScientific). To quantify protein band signals, X-ray films were scanned with a digital scanner and processed using Fiji ImageJ.

### Blood Hematological and Biochemical Analyses

For experiments in which mice were fed a high-fat diet, blood was obtained after fasting with *ad lib* in drinking water for 12-14 h to assess hematological and biochemical parameters. We obtained 300-500 µl of blood via left ventricular puncture from anesthetized mice. Blood was collected in tubes containing 20 µl of 0.5 M EDTA and 25 µl used within 2-3 h to obtain absolute monocyte counts. The remainder of the blood was centrifuged at 4°C at 1,900g for 10 min and the separated plasma was stored at -80°C until analysis. Plasma cholesterol and triglyceride concentrations were determined using an Element DC5X^TM^ Chemistry Analyzer (Heska) at the Institute of Comparative Medicine, Columbia University Irving Medical Center.

### Atherosclerotic Lesion and Vascular Tissue Analysis

We performed morphometric atherosclerotic lesion analysis as described previously.^37^ In brief, sections were stained with H&E and analysis performed by an experienced observer blinded to genotype. Total intimal lesion area from the internal elastic lamina to the lumen and acellular/anuclear areas (negative for hematoxylin-positive nuclei) per cross section were quantified by taking the mean of 6 sections spaced 30 µm apart beginning at the base of the aortic root. The necrotic core was defined as a clear area that was not stained by H&E. Images were viewed and captured with a Nikon Labophot 2 microscope equipped with an Olympus DP25 color digital camera attached to an imaging system with Image-Pro-Plus software (Media Cybernetics). An experienced observer blinded to genotype measured nuclei per area in aortic media, aortic media thickness, and adventitia thickness in H&E-stained sections. The density of nuclei (nuclei per mm^2^) in the aortic media was counted in one section per animal from each of the following four aortic regions: proximal ascending, mid arch, proximal descending, and lower descending. The thickness of the aortic media and adventitia in each animal was measured from the same four different regions of the aorta using Fiji ImageJ software. The means of three measurements in one section from each region were used for statistical analysis.

### Statistics

Statistical analyses were performed using GraphPad Prism 5. We used one-way ANOVA with Tukey’s multiple comparison test for comparisons between more than two groups. To assess statistical significance between two groups, we performed unpaired student’s two-tailed *t*-tests using Excel 2019 (Microsoft). Findings with P-value ≤0.05 were considered statistically significant.

## RESULTS

### Generating and Characterizing Hyperlipidemic *Ldlr*^−/−^ mice that express only prelamin A

To assess the effects of prelamin A on atherosclerosis, we generated *Lmna*^L648R/L648R^;*Ldlr*^−/−^ mice. We crossed *Lmna*^+/L648R^;*Ldlr*^+/+^ mice to *Lmna*^+/+^;*Ldlr*^−/−^ mice to generate *Lmna*^+/L648R^;*Ldlr*^+/-^ mice, which then were intercrossed to obtain *Lmna*^L648R/L648R^;*Ldlr*^−/−^ offspring. The intercrosses also generated *Lmna*^+/+^*;Ldlr*^+/+^, *Lmna*^L648R/L648R^*;Ldlr*^+/+^, and *Lmna*^+/+^;*Ldlr*^−/−^ offspring for comparisons. We restricted our analyses to male mice to avoid potential complications of sex differences that have been inconsistently reported in *Ldlr*^−/−^ mice.^38^

*Lmna*^L648R/L648R^*;Ldlr*^+/+^ and *Lmna*^L648R/L648R^*;Ldlr*^−/−^ mice expressed only prelamin A and no mature lamin A in livers and aortas, and *Lmna*^+/+^*;Ldlr*^+/+^ and *Lmna*^+/+^*;Ldlr*^−/−^ mice expressed only mature lamin A and no prelamin A (Figure 1A). There appears to be relatively more lamin C in *Lmna*^L648R/L648R^ mice compared to *Lmna*^+/+^ mice, but this is independent of the *Ldlr* allele. After eating a high-fat diet from 16 to 28 weeks of age, mice of all four genotypes had increases in body mass; those that were *Lmna*^L648R/L648R^ had slightly lower body masses at 28 weeks but the differences were not statistically significant (Figure 1B). After 12 weeks on a high-fat diet, *Lmna*^+/+^;*Ldlr*^−/−^ and *Lmna*^L648R/L648R^;*Ldlr*^−/−^ mice each had mean plasma cholesterol concentrations of approximately 1,300 mg/dl, as compared to approximately 100 mg/dl in *Ldlr*^+/+^ mice, and these values were not significantly different between the two different mouse groups with *Ldlr*^−/−^ genotype (Figure 1C). Mean plasma triglyceride concentrations were significantly increased in *Ldlr*^−/−^ mice compared to *Ldlr*^+/+^ mice but, also not significantly different between the two mouse groups with *Ldlr*^−/−^ genotype (Figure 1D). *Ldlr*^−/−^ mice had more severe hepatic steatosis than *Ldlr*^+/+^ mice as demonstrated on H&E-stained sections of livers, but the extent was similar in both mouse groups with *Ldlr*^−/−^ genotype (Figure 1E). There was no difference in the mean absolute blood monocyte between the genotypes (Figure S1). All the differences between *Ldlr^−^*^/-^and *Ldlr*^+/+^ mice are as expected and the absence of differences between the *Lmna*^+/+^ and *Lmna*^L648R/L648R^ mouse groups is a meaningful negative finding.

**Figure 1.**
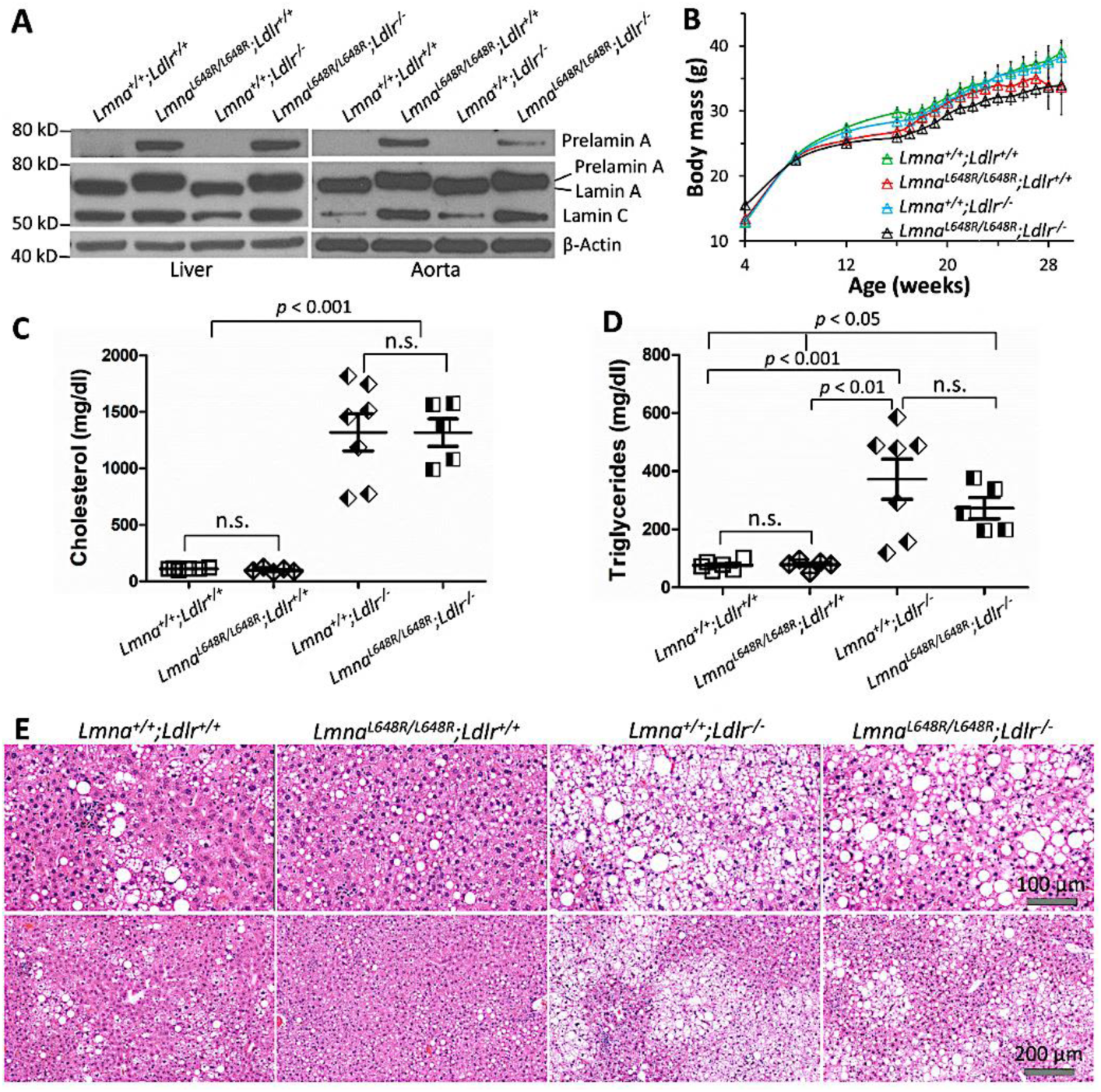
Control and hyperlipidemic mice that express only prelamin A or mature lamin A. **A,** representative immunoblots of proteins extracted from livers and aortas of mice of each indicated genotype probed with antibodies against prelamin A (top row), lamin A/C (middle row), and β-actin (bottom row). **B,** body mass versus age of male mice of each indicated genotype fed a high-fat diet from 16 to 28 weeks of age. Values are means ± SEM (n = 5-7). **C,** plasma cholesterol concentrations at 28 weeks of age. Each symbol is value for an individual mouse, and bars indicate means ± SEM (n =5-7). **D,** plasma triglyceride concentrations at 28 weeks of age. Each symbol is value for an individual mouse, and bars indicate means ± SEM (n = 5-7). **E,** Representative H&E-stained sections of livers at 28 weeks of age; fat vacuoles appear as empty white spaces within hepatocytes.

### Effects of Prelamin A on Aortic Atherosclerotic Lesions and Smooth Muscle in Hyperlipidemic Mice

After 12 weeks on a high-fat diet (from ages 16-28), both *Ldlr*^−/−^ mouse groups developed aortic root atherosclerotic plaques, whereas the aortas of *Ldlr*^+/+^ mouse groups were normal (Figure 2A). Notably, however, quantitative morphometric analysis of the plaques showed no difference between *Lmna*^+/+^;*Ldlr*^−/−^ mice that express only mature lamin A and *Lmna*^L648R/L648R^;*Ldlr*^−/−^ mice that express only prelamin A (Figure 2B). Similarly, plaque core necrotic areas were not different in *Lmna*^+/+^;*Ldlr*^−/−^ and *Lmna*^L648R/L648R^;*Ldlr*^−/−^ mice (Figure 2C).

**Figure 2.**
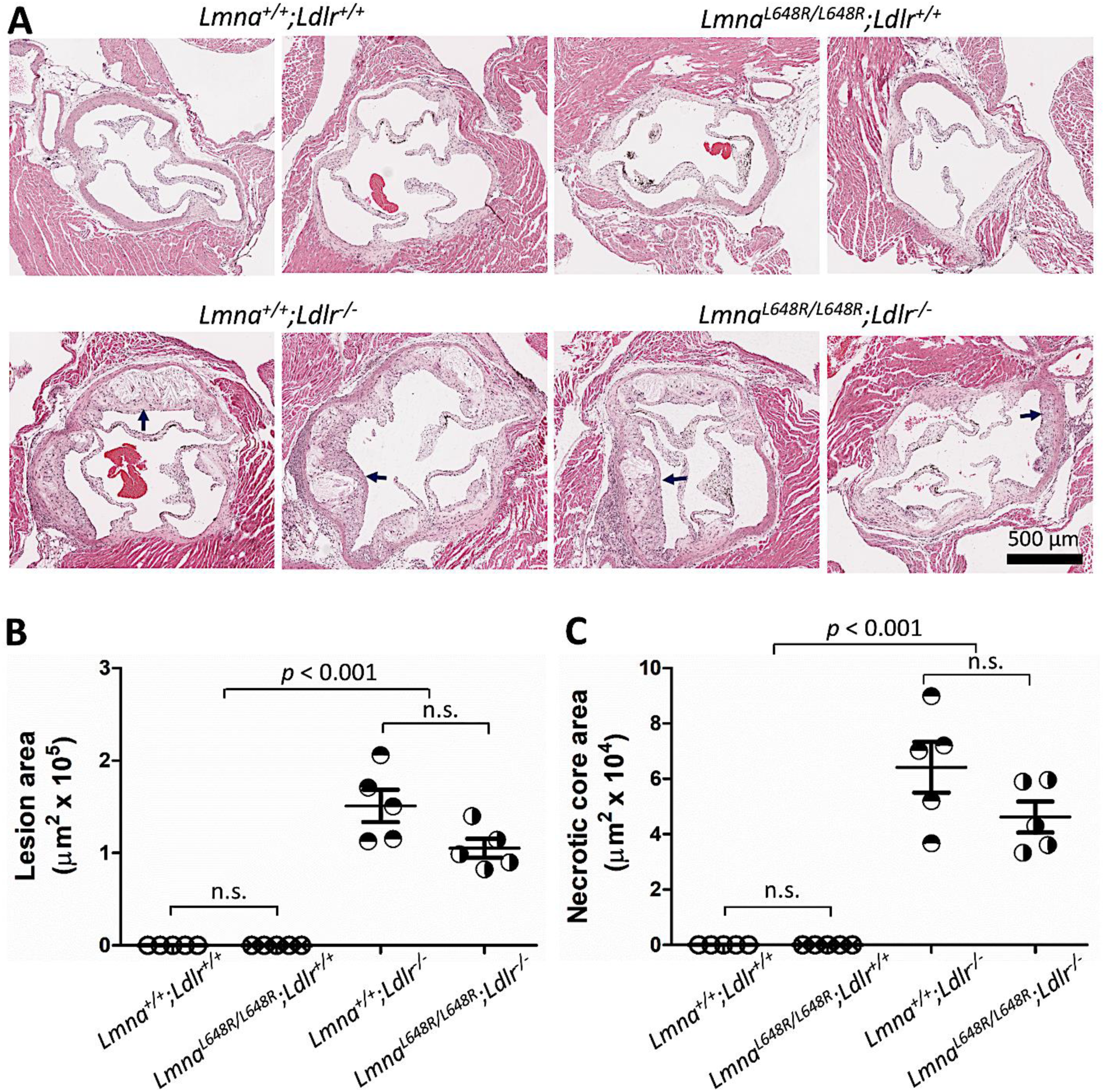
Aortic root atherosclerotic lesion and necrotic core areas in control and hyperlipidemic mice expressing only prelamin A or mature lamin A. **A,** representative H&E-stained sections of aortic roots of 28-week-old male mice of each indicated genotype fed a high-fat diet from 16 to 28 weeks of age. Arrows indicate representative atherosclerotic plaques. **B,** quantification of lesion area in aortic root sections. **C,** quantification of necrotic core area in aortic root sections. In B and C, each symbol is value for an individual mouse, and bars indicate means ± SEM (n = 5).

Severe loss of aortic VSMCs and a significantly increased adventitia to media thickness ratio are predominant features in humans and mice expressing progerin, the truncated prelamin A variant in HGPS.^10–18^ We therefore assessed media vascular smooth muscle in the aortas of atherosclerotic mice expressing prelamin A. In H&E-stained sections from mice of all genotypes, the media smooth muscle appeared normal in the proximal ascending aorta and mid aortic arch, even near atherosclerotic plaques in the LDL receptor-deficient mice, and in the proximal descending aorta and lower descending aorta (Figure 3A). The adventitia appeared normal on trichrome-stained sections of aortas from the mice of the four different genotypes (Figure S2). The number of nuclei per area of media in different portions of the aorta was not different in mice of these genotypes, indicating no detectable loss of VSMCs (Figure 3B). The ratio of adventitia to media thickness was also the same in different portions of the aorta in mice of all the genotypes (Figure 3C). In summary, expression of prelamin A had no discernible impact on aortic atherosclerotic plaques, the number of media VSMCs, or the adventitia in hyperlipidemic 28-week-old mice.

**Figure 3.**
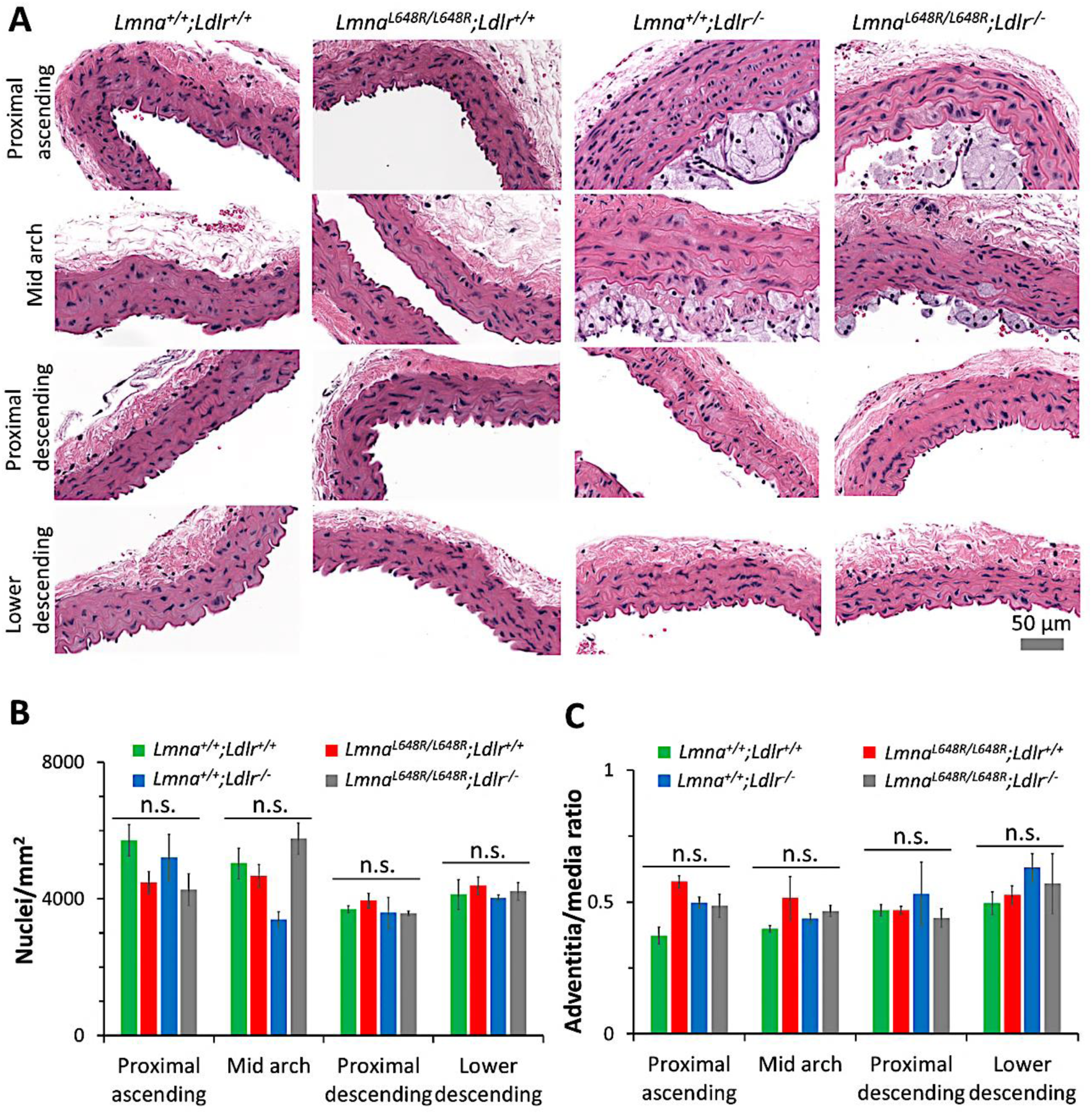
Aortic VSMCs per area and adventitial/media ratios in control and hyperlipidemic mice expressing only prelamin A or mature lamin A. **A,** representative H&E-stained sections from indicated regions of the aorta of 28-week-old male mice of each indicated genotype fed a high-fat diet from 16 to 28 weeks of age. See Figure S2 for trichrome-stained sections. **B,** quantification of number of nuclei per area in the media in indicated regions of the aorta. **C,** adventitial/media thickness ratios in indicated regions of the aorta. In B and C, values are means ± SEM (n = 3).

### Effects of Prelamin A on Aortic Smooth Muscle in Aged Mice

*Zmpste24*^−/−^ mice that express only prelamin A (no mature lamin A) do not have loss of VSMCs at 21 weeks of age.^30^ However, prelamin A has been reported to accumulate in carotid and aortic media of aged but not young humans and in atherosclerotic lesions and was proposed to play a role in vascular aging.^33,34^ We therefore used *Lmna*^L648R/L648R^ mice to determine if prelamin A has any effects on aortic smooth muscle in older mice at 52 weeks of age.

We fed *Lmna*^+/+^*;Ldlr*^+/+^, *Lmna*^L648R/L648R^*;Ldlr*^+/+^, *Lmna*^+/+^;*Ldlr*^−/−^ and *Lmna*^L648R/L648R^*;Ldlr*^−/−^ mice a regular chow diet from after weaning at 3 weeks to 52 weeks of age. At 52 weeks of age, we isolated aortas from male mice for analysis. Consistent with our previously reported findings,^31^ *Lmna*^L648R/L648R^ mice fed a regular chow diet had lower body masses than *Lmna*^+/+^ mice starting at approximately 10 weeks of age, regardless of whether they were LDL receptor-proficient or LDL receptor-deficient (Figure S3). In H&E-stained sections, the media smooth muscle appeared normal in the proximal ascending aorta, mid aortic arch, proximal descending aorta and lower descending aorta of mice of all these genotypes, including *Lmna*^L648R/L648R^;*Ldlr*^+/+^ and *Lmna*^L648R/L648R^;*Ldlr*^−/−^ (Figure 4A). The adventitia also appeared normal on trichrome-stained sections of aortas from the mice of the four different genotypes (Figure S4). The number of nuclei per area of media in different portions of the aorta was not different in mice of these genotypes, indicating no detectable loss of VSMCs (Figure 4B). The ratio of adventitia to media thickness was also the same in different portions of the aorta in mice of all genotypes (Figure 4C). These results showed that expression of only prelamin A did not cause loss of media VSMCs or adventitial thickening in mice at 52 weeks of age.

**Figure 4.**
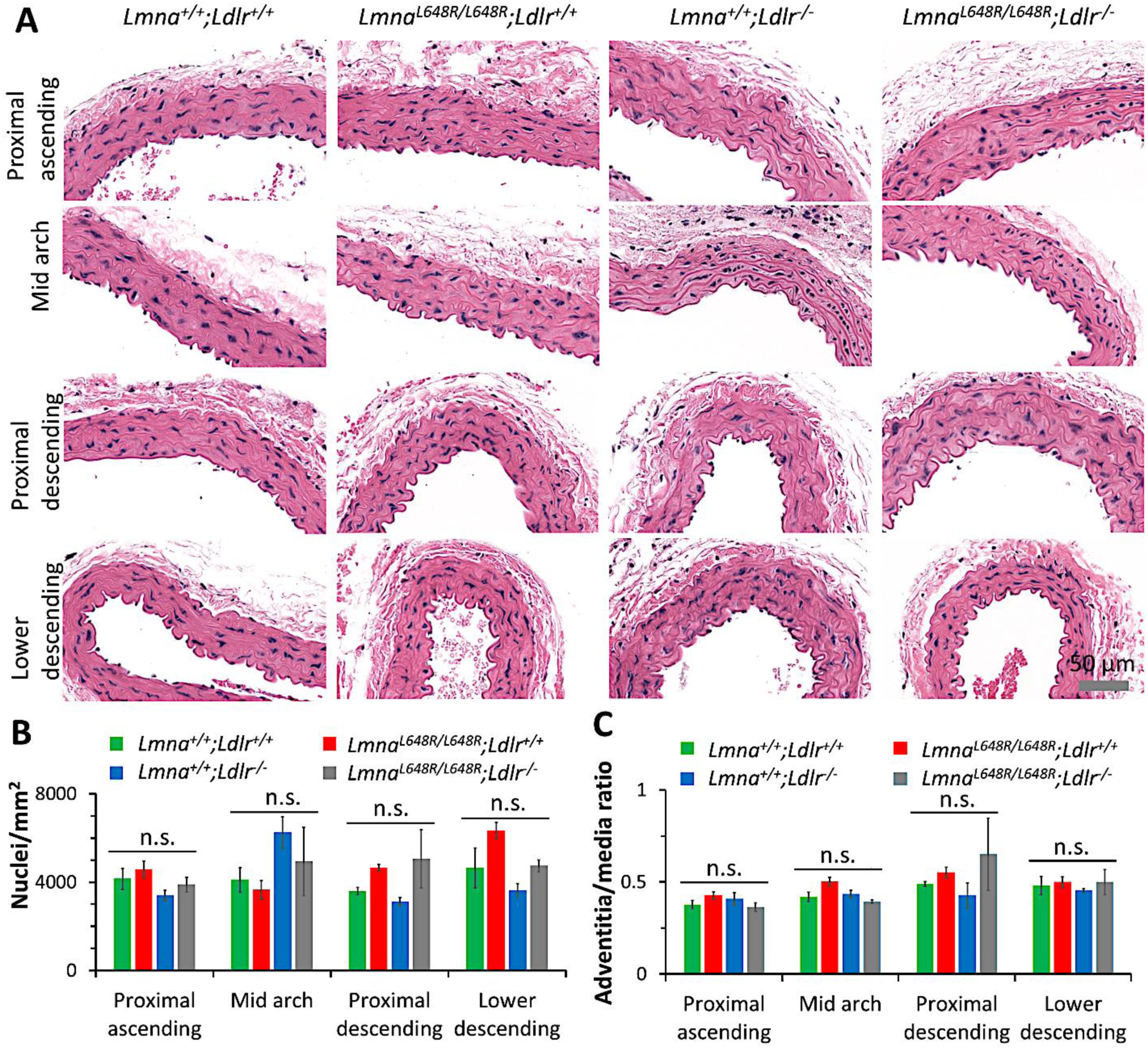
Aortic VSMCs per area and adventitial/media ratios in 52-week-old mice expressing only prelamin A or mature lamin A. **A,** representative H&E-stained sections from indicated regions of the aorta of 52-week-old male mice of each indicated genotype. See Figure S3 for trichrome-stained sections. **B,** quantification of number of nuclei per area in the media in indicated regions of the aorta. **C,** adventitial/media thickness ratios in indicated regions of the aorta. Values in B and C are means ± SEM (n = 3).

### Prelamin A Does Not Accumulate or Reduce AKT Activity in Aortas of *Lmna*^L648R/L648R^ Mic

Kim et al.^30^ reported that HGPS mice have decreased AKT activity in the aorta, which is responsible for decreased turnover of progerin and its accumulation with age; whereas in *Zmpste24*^−/−^ mice, aortic AKT activity was normal and prelamin A did not accumulate. We therefore determined if prelamin A accumulated with age in the aortic tissue of *Lmna*^L648R/L648R^ mice. We performed immunoblotting to measure prelamin A, lamin A, and lamin C in protein extracts from proximal descending aortas from 28-week-old and 52-week-old *Lmna*^+/+^*;Ldlr*^+/+^, *Lmna*^L648R/L648R^*;Ldlr*^+/+^, *Lmna*^+/+^;*Ldlr*^−/−^, and *Lmna*^L648R/L648R^*;Ldlr*^−/−^ mice (Figure 5A). There was no significant difference in lamin A or prelamin A expression levels at 28 weeks or 52 weeks in aortas from *Lmna*^+/+^*;Ldlr*^+/+^, *Lmna*^L648R/L648R^*;Ldlr*^+/+^, *Lmna*^+/+^;*Ldlr*^−/−^ and *Lmna*^L648R/L648R^*;Ldlr*^−/−^ mice (Figure 5B). Homozygosity for the *Lmna* L648R mutation did not therefore lead to increased expression or accumulation of the pathogenic prelamin A variant with age.

**Figure 5.**
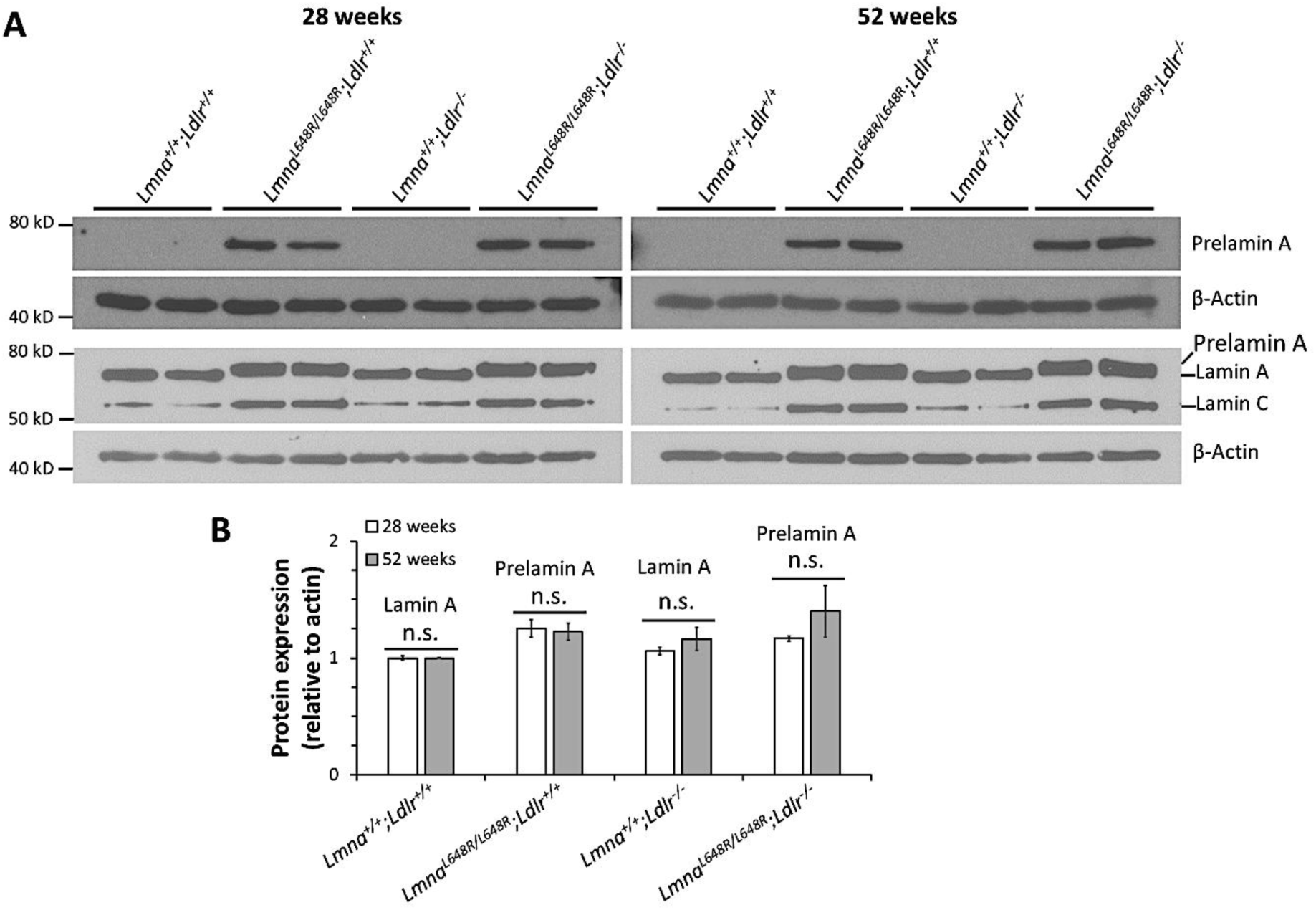
Prelamin A does not accumulate with age in aortas of *Lmna*^L648R/L648R^ mice. **A,** representative immunoblots of proteins extracted from aortas of 2 representative mice of each indicated genotype at 28 weeks of age and 52 weeks of age, probed with antibodies against prelamin A or β-actin (first and second rows) or against lamin A/C or β-actin (third and fourth rows). Two representative samples are shown for each genotype. **B,** quantification of lamin A or prelamin A in aortas of 28-week-old and 52-week-old *Lmna*^+/+^;*Ldlr*^+/+^, *Lmna*^648R/L648R^;*Ldlr*^+/+^, *Lmna*^+/+^;*Ldlr*^−/−^, and *Lmna*^L648R/L648R^;*Ldlr*^−/−^ mice. Values are means ± SEM (n = 3).

We next measured AKT activity, which has been shown to be reduced in aortas of HGPS but not *Zmpste24*^−/−^ mice^30^. We performed immunoblotting to measure AKT, pAKT (active), and the pAKT/AKT ratio in aortas from 28-week-old and 52-week-old *Lmna*^+/+^*;Ldlr*^+/+^, *Lmna*^L648R/L648R^*;Ldlr*^+/+^, *Lmna*^+/+^;*Ldlr*^−/−^, and *Lmna*^L648R/L648R^*;Ldlr*^−/−^ mice (Figure 6A). There were no differences in AKT (Figure 6B), pAKT (Figure 6C), or the pAKT/AKT ratio (Figure 6D) between these genotypes and young and aged mice. Hence, unlike what has been reported for progerin, expression of prelamin A does not reduce aortic AKT activity.

**Figure 6.**
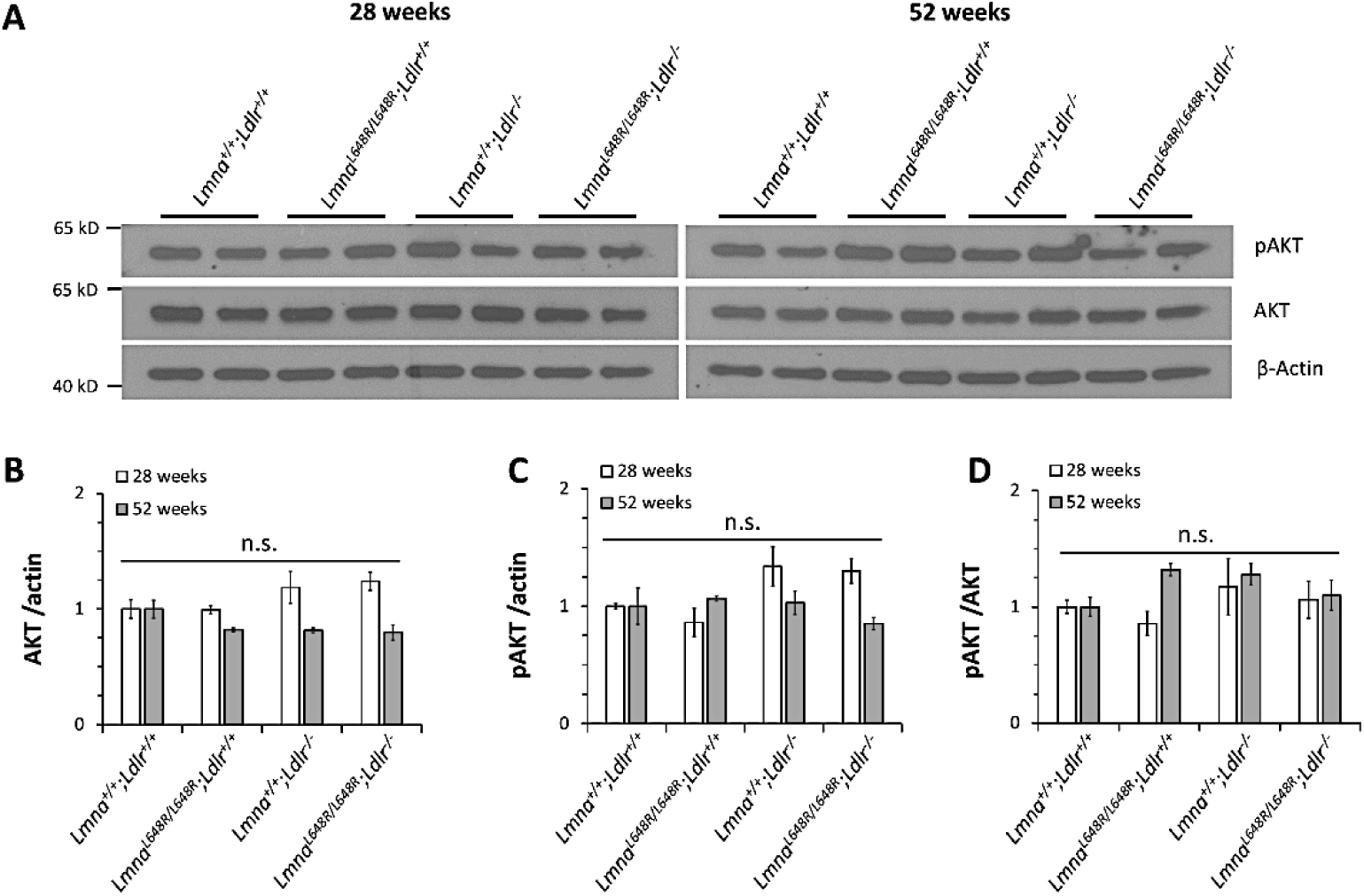
AKT activity does not change with age in aortas of *Lmna*^L648R/L648R^ mice. **A,** representative immunoblots of proteins extracted from aortas of mice of each indicated genotype at 28 weeks of age and 52 weeks of age probed with antibodies against pAKT, AKT and β-actin. Two representative samples are shown for each genotype. **B,** quantification of AKT in aortas of 28-week-old and 52-week-old mice of each indicated genotype. **C,** quantification of pAKT in aortas of 28-week-old and 52-week-old mice of each indicated genotype. **D,** pAKT/AKT in aortas of 28-week-old and 52-week-old mice of each indicated genotype. Values in B, C, and D are means ± SEM (n = 3).

## DISCUSSION

### Does Prelamin A Play a Role in Atherosclerosis or Vascular Aging?

We have shown that accumulation of prelamin A in *Lmna*^L648R/L648R^ mice does not accelerate or worsen atherosclerosis in hyperlipidemic mice. Aberrant accumulation of prelamin A also does not cause the typical aortic vascular pathology, including loss of medial VSCMs, that occurs with accumulation of progerin, the prelamin A variant in HGPS. These findings challenge the relevance of previous publications suggesting that prelamin A may accelerate smooth muscle cell senescence and is a novel biomarker of human vascular aging. Ragnauth et al.^33^ reported that arterial VSMCs isolated from aged individuals accumulated prelamin A upon passaging and that in human arteries, prelamin A expression was prevalent in medial VSMCs from aged individuals and in atherosclerotic lesions. They further reported that prelamin A accumulation correlated with decreased expression of ZMPSTE24 and that overexpression of prelamin A accelerated senescence of VSMCs. Liu et al.^34^ reported that prelamin A accumulated in arteries from young children on dialysis who develop hallmarks of premature vascular aging, including loss of VSMCs and medial calcifications. These authors further presented findings suggesting that prelamin A promotes calcification and aging of VSMCs by inducing persistent activation of the DNA damage response and senescence-associated secretory phenotype.

However, the present study as well as work by others discussed below contradict or are not aligned with the findings discussed above. We report here that in *Lmna*^L648R/L648R^ mice, which express only farnesylated prelamin A and no mature lamin A, all cells in which lamin A is normally expressed have no vascular pathology. Nor has vascular pathology been noted in clinical reports of patients with MAD-B with mutations in *ZMPSTE24* whose cells also express farnesylated prelamin A.^19–23^ There is one case report of a patient who underwent coronary angioplasty for chest pain at age 31 years, but he had chronic renal failure and severe hypertension.^39^ Furthermore, *Zmpste24*^−/−^ mice do not develop aortic pathology in their short lifetimes. Kim et al.^30^ recently showed that genetically-mediated excess expression of prelamin A in VSMCs – leading to total aortic amounts approximately 1.5 times greater than in *Zmpste24*^−/−^ mice – did lead to loss of VSMCs. However, the amount of prelamin A in arterial VSMCs from aged or atherosclerotic humans is unlikely to reach the amount observed in aortas of *Lmna*^L648R/L648R^ mice or 1.5 times the amount in aortas of *Zmpste24*^−/−^ mice. Therefore, even if prelamin A accumulates in some arterial VSMCs with human aging, it is extremely unlikely to contribute to vascular pathology, given the lack of any vascular abnormalities reported here for both hyperlipidemic and old *Lmna*^L648R/L648R^ mice.

### Differences Between Prelamin A and Progerin in Arterial Vascular Pathology

The aortic pathology in HGPS mice that express progerin versus *Lmna*^L648R/L648R^ mice that express prelamin A differs dramatically. Mice that express progerin, as well as children with HGPS, have a severe loss of VSMCs in the aortic media with adventitial fibrosis and thickening.^10–16^ We did not observe any of these changes in aortas of *Lmna*^L648R/L648R^ mice, even up to 52 weeks of age. *Zmpste24*^−/−^ mice that express prelamin A similarly do not have loss of VSMCs.^30^ Why does progerin have such dramatic effects on VSMCs whereas full-length prelamin A does not, even though both are permanently farnesylated forms of prelamin A?

A reasonable hypothesis is that the internal deletion of 50 amino acids in progerin, as compared to full-length prelamin A, may alter its half-life, function, or both in the aorta. Indeed, Kim et al.^30^ have shown that progerin and prelamin A are equally “toxic” in cultured VSCMs engineered to express an equivalent amount of these proteins. However, the picture was dramatically different when they compared aortas from aging HGPS and *Zmpste24*^−/−^ mice; progerin levels increased dramatically in the former, but not prelamin A in the latter. Their results suggested that the high level of progerin that accumulates in the aortas of HGPS mice compared to the lower levels of prelamin A in the aortas of *Zmpste24*^−/−^ mice promotes vascular pathology. Kim et al.^30^ posited that progerin is stable, with a slower turnover rate than prelamin A, in the context of the aorta. This is consistent with the finding that HGPS mice have an age-related increase in progerin in the aorta in the setting of decreasing transcript levels.^40^ Independent proteomic analysis of metabolically-labelled proteins has shown that progerin has a relatively slow turnover rate in aortas of HGPS mice.^41^ Furthermore, Kim et al.^30^ showed that raising the level of prelamin A in *Zmpste24*^−/−^ mouse aortas through genetic engineering resulted in vascular pathology, supporting the notion that it is the accumulation of progerin and that causes pathology. AKT-catalyzed phosphorylation of prelamin A has been shown to trigger its degradation.^42^ Kim et al.^30^ showed reduced AKT activity and AKT-catalyzed phosphorylation of progerin in aortas of HGPS mice but no reduction in AKT activity in aortas of *Zmpste24*^−/−^ mice. We similarly showed no reduction in AKT activity in aortas of *Lmna*^L648R/L648R^, further supporting the hypothesis that progerin but not prelamin A reduces AKT activity.

Another hypothesis regarding the different effects of progerin and prelamin A on the aortic vasculature is the difference between the proteins themselves. Most notably, progerin has an internal in-frame deletion of 50 amino acids. While the loss of these amino acids may enhance the “toxicity” of progerin relative to prelamin A, it appears that the accumulation of a high level of progerin that is the key to vascular pathology.^30^ In any case, there is little to no information on the function that the region deleted from progerin has in prelamin A.

### Limitations

Our study has several limitations. *Lmna*^L648R/L648R^ mice express prelamin A with a single amino acid substitution. It is possible that the L648R variant is less “toxic” than native prelamin A. However, this explanation is difficult to reconcile with the wealth of data suggesting that it is the farnesyl and carboxymethyl moieties of prelamin A that are primarily responsible for its adverse effects.^5,6^ It is possible that prelamin A L648R is a weaker substrate for protein farnesyltransferase than native prelamin A, leading to less farnesylation, which could render it less “toxic.” However, this is unlikely, since a 4-residue carboxyl-terminal CAAX motif – or its 3-residue or 5-residue variant – is sufficient for recognition by protein farnesyltransferase^43,44^ whereas L648R lies 14 residues upstream of the CAAX motif in prelamin A. Additionally, consistent with our data for *Lmna*^L648R/L648R^ mice, *Zmpste2*4^−/−^ mice and MAD-B patients that express native prelamin A also do not develop vascular pathology. ^19–22,26,30^

We studied aortic pathology in only male mice to avoid potential complications of sex differences that have been inconsistently reported in *Ldlr*^−/−^ mice.^38^ Hamczyk et al.^17^ similarily restricted their analysis of atherosclerosis-prone model of HGPS generated by crossing *Apoe*^−/−^ with *Lmna*^G609G/G609G^ to male mice. In another model generated by crossing *Ldlr*^−/−^ mice to *Lmna*^G609G/G609G^, the same group studied both sexes and did not report differences in aortic pathlogy.^18^ Other studies examined aortic pathologh in *Zmpste24*^−/−^ or *Lmna*^G609G/G609G^ mice ether analyzed male and female mice together or did not specify the sex used.^13,15,16,30,40^ We previously reported that female *Lmna*^L648R/L648R^ mice had longer lifespans than their male counterparts.^31^ Therefore, we do not believe that an analysis of female mice in the current study would have shown more severe effects of prelamin A on aortic pathology.

Finally, while widely used to study human atherosclerosis, there are limitations of mouse models.^45^ In *Ldlr*^−/−^ or *Apoe*^−/−^ mice fed a high-fat diet, atherosclerotic lesions primarily develop in the aorta and carotid arteries, not the coronary arteries. Atherosclerotic lesions in mice rarely rupture, which is often the cause of myocardial infarction in humans. Therefore, it is possible, although not yet demonstrated, that small amounts of prelamin A accumulation in human atherosclerotic coronary arteries may contribute to pathology.

## Non-standard Abbreviations

CAAX: cysteine-aliphatic-aliphatic-any amino acid
H&E: hematoxylin and eosin
HGPS: Hutchinson-Gilford progeria syndrome
MAD-B: mandibuloacral dysplasia with type B lipodystrophy
VSMCs: vascular smooth muscle cells

## Acknowledgments

We thank Dr. Ira Tabas (Columbia University) for assistance analyzing atherosclerotic plaques and Dr. Alan R. Tall (Columbia University) for providing *Ldlr*^−/−^ mice.

